# Age related disorder in DNA methylation patterns underlies epigenetic clock signals, but displays distinct responses to epigenetic rejuvenation events

**DOI:** 10.1101/2022.12.01.518745

**Authors:** Emily M. Bertucci-Richter, Ethan P. Shealy, Benjamin B. Parrott

## Abstract

Changes in DNA methylation with age are observed across the tree of life. The stereotypical nature of these changes can be modeled to produce epigenetic clocks capable of predicting biological age with unprecedented accuracy. Despite the predictive ability of epigenetic clocks, the underlying processes that produce clock signals are not resolved but are hypothesized to be rooted in stochastic processes leading to an erosion in the epigenetic landscape. Here, we test this hypothesis using a novel computational approach for measuring disorder in DNA methylation patterns across the epigenome. We find that loci comprising conventional epigenetic clocks are enriched in regions that both accumulate and lose disorder with age, suggesting a direct link between DNA methylation disorder and epigenetic clock signals. Across the murine lifespan, disorder accumulates in Polycomb Repressive Complex 2 target genes and decreases in CTCF and transcription factor binding sites, resulting in genomic hotspots of age-related epigenetic disorder. We further investigate the connections between age-related changes in disorder and epigenetic clock signals by comparing the influences of development, lifespan interventions, and cellular dedifferentiation using a series of newly developed epigenetic clocks based on regional disorder, average methylation states, and commonly used measures of entropy. We identify both common responses as well as critical differences between canonical epigenetic clocks and those based on regional disorder, demonstrating a fundamental decoupling of epigenetic aging processes. Collectively, this work identifies key linkages between epigenetic disorder and epigenetic clock signals, and simultaneously demonstrates the multifaceted nature of epigenetic aging in which stochastic processes occurring at non-random loci produce predictable outcomes.

## Introduction

Changes in DNA methylation with age, or “epigenetic aging”, are widely observed across the tree of life. Age-associated DNA methylation patterns manifest as two general phenomena; one leading to stereotypical shifts in mean methylation levels at individual cytosines that can be modeled to predict individual age with high accuracy [1], and the other leading to increased variability or “disorder” in DNA methylation states due to the erosion of the epigenetic landscape [2–4]. These phenomena are hypothesized to be linked as average methylation values of individual cytosines are reported to drift from hyper- or hypo-methylated (e.g., ≥ 80%, ≤ 20%) states to more intermediate levels (e.g., 20%- 80%) with age [5]. However, the extent to which age-associated changes to the DNA methylome reflect distinct or similar underlying processes remains unresolved.

Over the last decade, dozens of epigenetic clocks have been developed for a range of taxonomic groups including humans [6], rodents [7], fish [8], birds [9], and trees [10]. Epigenetic clocks are typically constructed as linear models that predict chronological age or age-related phenotypes using mean methylation levels from a relatively small number of individual cytosines. The rate of epigenetic aging, measured as the discrepancy between chronological age and epigenetic age estimates, is associated with environmental conditions [6], life history traits (e.g., age at first menarche [11] and menopause [12]), and has become a widely used indicator of biological age and attendant disease risk [1,13,14]. More recently, epigenetic clocks have been applied to understanding epigenetic rejuvenation events occurring either naturally during early embryonic development or as a consequence of cellular reprogramming. Whereas epigenetic age estimates of induced pluripotent stem cells (iPSCs) are typically reset to zero [6], transient treatments with Yamanaka factors that do not fully induce dedifferentiation also reduce epigenetic age estimates and have been recognized as a promising anti-aging therapeutic avenue [15]. Kerepesi, et al. (2021) have also reported a period of epigenetic rejuvenation occurring during early development in which epigenetic age estimates decrease after conception until reaching a “ground zero” state coinciding with gastrulation [16]. Yet, age estimates derived from epigenetic clocks may not fully capture other facets of epigenetic aging, and here, we integrate multiple measures of age-associated DNA methylation patterns to examine these phenomena more broadly.

The mechanistic underpinnings of epigenetic clock signals are still unclear, but with millions of CpG dinucleotides in the genome [17], and minimal overlap of individual CpGs included across different epigenetic clocks [18], epigenetic aging is suggested to be the product of a more general epigenetic maintenance system [1]. Commonly referred to as epigenetic “drift”, the failure of this maintenance system has many references in the recent literature [2,19–23]. Yet, despite an abundance of reports examining age-related epigenetic drift [19,22,24], a consensus definition is lacking, with studies often defining drift to mirror the analytical approach employed [25]. For example, “epiallele frequency” [26], “discordance” [27], “disorder” [28], “entropy” [29], and “heterogeneity” [30] have all been used to assess epigenetic drift and reflect different analytical approaches. Perhaps the most inclusive definition of epigenetic drift is a change in the status of DNA methylation over time [22,24]. Yet, according to this definition, even programmed changes which guide developmental processes could be considered epigenetic drift, and it is likely more useful to define epigenetic drift as a stochastic, rather than a deterministic change in methylation states. One popular approach for assessing stochastic changes in methylation is using Shannon’s Entropy [31]. Originating in information theory, this metric measures the amount of uncertainty in an occurrence or event. However, when applied to DNA methylation, Shannon’s Entropy simply reflects average methylation values (whether genome wide or at a specific CpG) and is also likely influenced by heterogeneity among cells. Heterogeneity of epigenetic patterning *within* cells requires analyzing single cells or in the case of bisulfite sequencing experiments, can be inferred from linked CpGs occurring on individual reads [28,32].

Herein, we apply novel read-based strategies to resolve age-associated epigenetic disorder across the mouse genome. By considering methylation states between individual CpGs and their immediate neighbors, we directly assess epigenetic disorder and investigate its relationship to epigenetic clock signals, embryonic development, lifespan interventions, and cellular reprogramming. Borrowing from the conceptual framework of Waddington’s epigenetic landscape, we hypothesize that low levels of epigenetic disorder characterize robust epigenomic states and that gains in disorder occurring with age lead to “erosion” of this landscape [2,33–36]. We find that approximately 30% of the genome is disproportionately affected by age-related epigenetic disorder. Loci which act as predictors in conventional epigenetic clocks based on mean methylation levels appear to be enriched in regions that both accumulate and lose disorder with age, suggesting a direct link between epigenetic disorder dynamics and clock signals. We subsequently develop a new class of epigenetic clocks based on regional disorder (RD) metrics and compare these signals with those produced using conventional epigenetic clocks and those based on entropy. Upon exploring the influences of development, lifespan interventions, and cellular dedifferentiation, we identify similarities as well as clear divergence between epigenetic clock signals based on either mean DNA methylation or regional DNA methylation disorder. Contrary to predictions based on prior studies, we find that disorder increases during early development and global levels of disorder are unaffected after cellular rejuvenation. Collectively, our findings suggest that DNA methylation disorder dynamics are a key contributor to epigenetic clock signals, yet also highlight a fundamental decoupling of disorder dynamics from canonical epigenetic aging that is likely to inform the potential of lifespan intervention strategies.

## Materials and Methods

### Data acquisition

Reduced representation bisulfite sequencing (RRBS) data from 255 mouse samples were acquired from NCBI’s Sequence Read Archive (Accession: PRJNA319643). Individuals ranged in age from 0.67 to 35 months, and represented both sexes, four strains (DW/J x C3H/HEJ)/F2, (C57BL/6J x BALB/cByJ)/F2, B6D2F1 and C57BL/6), and two diets (standard and caloric restriction). This dataset included methylomes from whole blood samples, induced pluripotent stem cells (iPSCs) derived from kidney (n = 3) and lung (n = 3), as well as the fibroblasts they were derived from (n = 3 lung, n = 3 kidney). Sample collection and library preparation methods are detailed in [37].

### Data processing

Raw sequence reads were trimmed of low-quality sequences using Trim Galore! (v0.6.5, options: --paired –rrbs –quality 25 –illumina). Trimmed reads were then aligned in paired end mode to a bisulfite index of the latest version of the mouse genome (GRCm39) using Bismark (v0.22.3), with mapping efficiency ranging from 54-70% among samples. Following alignment, reads were sorted by genomic coordinate, and converted to human readable SAM files using the Samtools (v1.10) functions ‘sort’ and ‘view’, respectively. The methylation call strings from each read were extracted using a custom R script. Reads with less than 2 CpGs were removed from the analysis. Each CpG within a methylation call string was then scored based on whether its methylation status matched the methylation status of its nearest neighbors. Because the first and last CpG on a string has only one nearest neighbor, the maximum disorder score is one (1), while each CpG in the internal part of a string has two nearest neighbors (one upstream and one downstream), giving a maximum disorder score of two (2).

### Calculation of disorder

The proportion of disordered neighbor pairs (PDN) was calculated on a per read basis by taking the proportion of neighbor pairs within the read that were disordered (i.e. methylation state differed) over the total number of neighbor pairs within the read. Practically, this was calculated as follows:

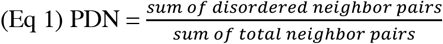

### Calculation of Regional Disorder and Methylation

Due to differences in coverage across individuals, we normalized our metric of disorder across 200 bp windows of the genome, subsequently referred to as regional disorder (RD; Figure 1A). To measure RD, we binned the genome into 200 bp windows using the Bedtools (v2.26.0) function ‘makewindows’ and used the Bedtools ‘map’ function to average the per-read PDN, methylation, and CpG density for all reads for which >51% of the read mapped to a specific window, preventing reads from being represented in more than one region. Regional methylation (RM) was calculated using the mean proportion of methylated cytosines within each region. Regions with less than five reads per sequencing run were excluded from analysis, and data from separate sequencing runs were merged together on a per individual basis using a weighted average based on the number of reads from each run. We then removed regions which were not present in at least 80% of all 255 samples.

**Figure 1.**
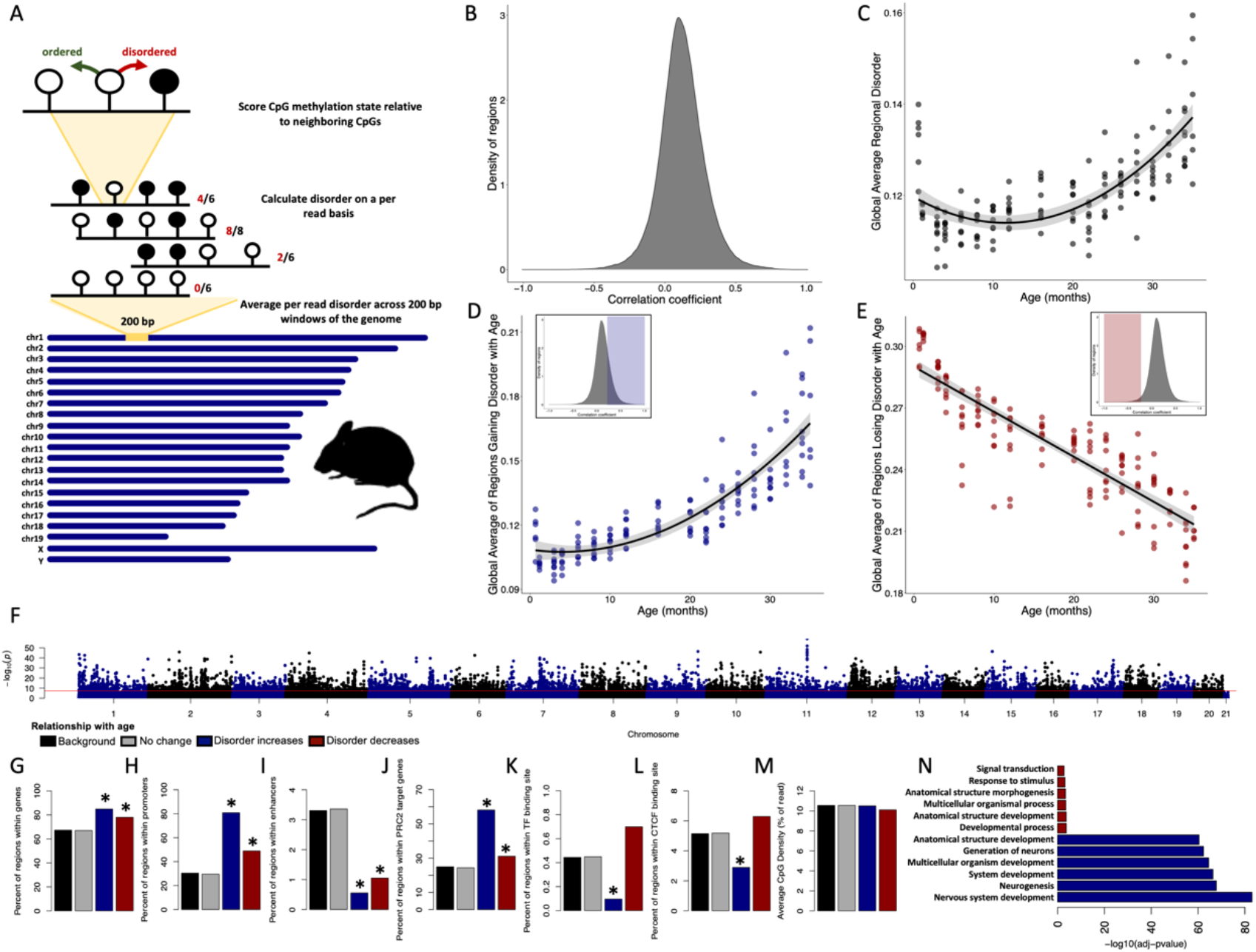
Epigenetic disorder increases across the murine lifespan. (A) Diagram of the approach for measuring regional disorder (RD). (B) Density of all genomic regions assessed with respect to their Spearman correlation coefficients between RD and age. (C) The relationship between global disorder and age in mice. (D) Average RD across all regions that gain disorder with age (correlation coefficient ≥ 0.25), or (E) lose disorder with age (correlation coefficient ≤ −0.25. (F) Manhattan plot of the distribution of FDR corrected p-values of the relationship between RD and age. Enrichment of age associated RD in genes (G), promoters (H), enhancers (I), PRC2 target genes (J), transcription factor binding sites (K), CTCF binding sites (L), and average CpG density (M). (N) The six most significant gene ontology biological processes (GO:BP) for regions gaining or losing disorder with age. Regions which gain disorder with age are shown in blue and regions which lose disorder with age are shown in red.

### Calculation of Regional Entropy

Regional entropy (RE) was calculated for each 200 bp window as follows:

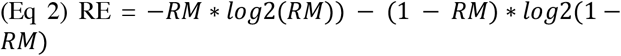

### Age-associated disorder, methylation, and entropy

To test if disorder increased with age, we selected a subset of whole blood methylomes from 153 male, C57BL/6 mice fed a standard diet, with individuals ranging from 0.67 to 35 months of age. Using this subset, we performed individual Spearman correlations between age and both RD and RM with a false discovery rate (FDR) correction for multiple comparisons using the corr.test function from the package psych in R [38]. Regions with a correlation coefficient ≥ 0.5 and an FDR corrected p-value < 0.05 were considered to gain disorder or methylation with age, and those with a correlation coefficient ≤ −0.5 and p-value < 0.05 were considered to lose disorder or methylation with age.

### Calculation of global disorder

For each sample we calculated global disorder using the mean RD values of those regions passing the filtering approach outlined above, which allowed us to directly compare disorder between individuals despite differences in coverage or depth of coverage across the genome. We also calculated global disorder using only regions which displayed any modest gain disorder with age (correlation coefficient ≥ 0.25; n = 45,668) and only regions which lost disorder with age (correlation coefficient ≤ −0.25; n = 3,789). We then modeled the relationships between all three global disorder metrics and age using the lme package in R, and age-adjusted global disorder was calculated using the residuals from the quadratic relationship between global disorder and age.

### Genomic enrichment of age-associated disorder

We then classified each 200 bp region by its genomic localization according to annotations of genes, introns, exons, CpG density, promoters, enhancers, transcription factor (TF) binding sites, CTCF binding sites, polycomb repressive complex 2 targets (PRC2), and Petkovich epigenetic clock sites [37]. Coordinates for genes, introns, and exons were used as listed by the most recent Refseq annotation of the mouse genome (GRCm39) with genes considered as the entire interval between transcription start and end coordinates. Coordinates of promoters, enhancers, TF binding sites and CTCF binding sites were determined using the Expression and Regulation annotation track from UCSC Genome Browser (GRCm39), CpG density was calculated using the average number of CpGs per informative read over the 100 possible CpGs per region, and PRC2 target genes were determined by previously published ChIP-seq data of PRC2 subunit binding in mouse ESCs [39], with any gene binding at least one PRC2 subunit being considered a PRC2 target. Coordinates from the Petkovich epigenetic clock [37] were translated to the current mouse genome annotation using NCBI’s coordinate remapping service. Overlap between the 200 bp regions and each genomic category (at least 1 bp) was determined using a custom R script. Genomic enrichment was determined using binomial tests using all other covered loci as a background.

### Gene Ontology

Genes in regions determined to gain or lose RD with age were split into lists and compared against the background (all represented genes) for gene ontology enrichment using gProfiler. Genes spanning multiple age-associated regions were only counted once per gene list.

### CpG Methylation

Merged alignment files for each sample were also used to produce CpG methylation matrices using Bioconductor’s MethylKit. Individual cytosines from opposite strands were merged into single CpGs (destrand = TRUE). Only CpGs which were covered at a depth of 10x reads across all 153 male, C57BL/6, standard diet samples were retained for further analysis.

### Clock optimization

To compare our measures of disorder with epigenetic aging, we developed four different epigenetic clocks based on RD, RM, RE, and CpG methylation as predictors of chronological age. We used the glmnet package in R to select predictors using elastic net regularized regression and a leave-one-out cross validation (LOOCV) approach to assess model performance. Alpha values for each model were set to 0.5 (true elastic net) and lambda was cross validated across all samples in the training set for each individual model. Age estimates from test samples (i.e., remaining individuals not used to train the model) were used to assess the error of the clocks. To assess robustness of individual predictor sites, we extracted predictors from each model and determined the proportion of the 153 data-type-specific clocks each was included in. The robustness of CpG clocks was assessed by assigning individual CpGs to their respective genomic region, with each region being counted only once per clock iteration (i.e. multiple clock sites per region were not multiply counted.) We then determined the overlap between selected clock regions between RM, RD, RE, and CpG clocks.

### Representative clock building

While LOOCV approaches provide a more inclusive estimate of predictive power, they do not provide a singular model appropriate for downstream applications. Thus, we constructed an additional set of clocks by randomly splitting samples into a training set (n = 14) and a test set (n = 39), which consisted of 2 or 3 individuals from each age class. We refer to these models as the “representative” clocks for each data type (Figure S1), and the same training and test set were used for every data type.

### Testing the effects of lifespan interventions

We then tested the effects of three lifespan interventions: caloric restriction beginning at 14-weeks of age, knock out of growth hormone receptor (GHR), and dwarfism using the representative clocks. The dataset consisted of 22 male and female individuals from mouse strain Snell Dwarf (DW/J x C3H/HEJ)/F2, split between Snell Dwarf (mutation in Pit-1 gene; n = 10) and their respective controls (“Snell Dwarf Control”; n = 12), 26 male and female individuals from strain (C57BL/6J x BALB/cByJ)/F2, split between GHR knock out (GHRKO, n = 11) and GHR wild type (GHR WT, n = 15), 22 male B6D2F1 mice, split between standard diet (n = 10) and caloric restriction (n = 12), and 20 male individuals from line C57BL/6 on a calorie restricted diet. Specific details of lifespan extending treatments can be found in Petkovich, et al. (2017) [40]. We calculated age adjusted global disorder, RD, RM, and RE as described above, and extracted CpG methylation information for each individual and then applied our representative epigenetic clocks from each data type to acquire epigenetic age estimates. Data from individuals experiencing lifespan interventions was handled exactly as described above and any missing predictor was assigned a zero value so as to be dropped from the model. Differences between treatments were determined using a one-way ANOVA.

### Disorder during de-differentiation and development

We analyzed DNA methylomes from mouse iPSCs and their respective lung (n = 6) or kidney (n = 6) fibroblast precursors. Specific details regarding the de-differentiation of fibroblasts can be found in Petkovich, et al. (2017) [40]. Datasets for the analysis of methylation dynamics across embryonic development were acquired from SRA accessions PRJNA150129 and PRJNA221793. Methods for sample preparation and sequencing in these datasets are detailed in Kerepesi, et al. (2021)[16] and Smith, et al. (2012) [41], respectively. Sample selection and filtering for loci comprising the Stubbs epigenetic clock [7] was modeled after the epigenetic clock methods in Kerepesi, et al. (2021)[16] to reproduce reported results with the traditional CpG-based approach. This included removal of samples retaining the polar bodies, as well as those derived from pre-fertilization gametes and ESCs. A similar sample-selection strategy was utilized for the region-based metrics, but the filtering strategy instead followed that outlined earlier in this paper for RD, RM, and RE. Overall, 36 samples were included in the window-based analyses, and 38 in the CpG-based Stubbs clock (due to differences in filtering requirements between the two approaches). Developmental stages represented in the ‘early’ developmental group ranged from zygote to ICM (approximated to 0.5-3.5 days after [16]), with the ‘late’ group consisting of embryonic and extraembryonic tissue from E6.5 and E7.5 embryos.

Age adjusted global disorder, and epigenetic age estimates for embryonic samples were calculated using representative RD, RM, RE clocks and the Stubbs CpG clock [7] as described for the lifespan intervention experiments. As data originated from two different datasets and consisted of different tissues than those used to train representative clocks, age adjusted global disorder and epigenetic age predictions were normalized within their respective datasets. Differences between the epigenetic ages of iPSCs were determined using a two-way ANOVA with cell type (iPSC or fibroblast) and tissue (kidney or lung) as predictors. To further investigate the role of disorder during development and de-differentiation, we performed two-tailed t-tests to determine differences in RD occurring after de-differentiation (fibroblast vs iPSC), or across development. For this analysis, we grouped both fibroblast types (kidney and lung) to compare against the iPSCs, as well as grouping the developmental datasets into early (E0.5-3.5) and late (E6.5-7.5) development. Given the especially low sample size for the iPSC dataset (n = 12), we also removed any regions with missing values. P-values from t-tests were corrected using FDR, via the function p.adjust in R. Significant differences between groups were determined by an adjusted p-value ≤ 0.05 and a mean difference in disorder between groups of at least |0.1|. Significant differences in disorder were then further characterized into regions which gained disorder during development or de-differentiation, and regions which lost disorder during development or de-differentiation. To determine the effect size of any given region on epigenetic age prediction, we took the mean difference between groups (either de-differentiation or development) at that region and multiplied it by the beta value for that region used in the RD epigenetic clock model. The effect size for each region was then normalized to the percent of the total effect size for the clock.

## Results

Disorder in DNA methylation patterns are strongly correlated with age on a regional and global scale. Of the 249,015 regions assessed, RD was significantly correlated with age in 76,353 regions (30.7%), with RD increasing with age in 70,094 genomic regions (91.8%; Figure 1B) and decreasing with age in 6,259 genomic regions (8.2%; Figure 1B). The average RD across all regions, or global disorder, increases with chronological age according to a quadratic relationship (R^2^ = 0.51, p < 2.2e-16; Figure 1C). Consistent with increases and decreases in RD being driven by distinct processes, regions experiencing increases in RD (cor ≥ 0.25) display a quadratic relationship to age (R^2^ = 0.74, p < 2.2e-16; Figure 1D), whereas regions experiencing decreasing RD (cor ≤ 0.25) display a linear relation with age (R^2^ = 0.77, p < 2.2e-16; Figure 1E).

With the exception of the Y chromosome, every chromosome incurs significant age-related accumulation of RD with “hot spots” of age-related disorder occurring across the genome (Figure 1F). Given that more than 30% of the genome experiences age-associated RD, only sites with a p-value ≤ 0.05 and a correlation coefficient greater ≥ 0.5 (n = 4149) or ≤ −0.5 (n = 286) were considered as age-associated for enrichment tests, with all other regions considered background (n = 244,580). Regions accumulating RD with age were significantly enriched in genes (p < 2.2e-16; Figure 1G) and promoters (p < 2.2e-16; Figure 1H), depleted in enhancers (p < 2.2e-16; Figure 1I), enriched in PRC2 target genes (p < 2.2e- 16; Figure 1J), and depleted in both transcription factor (p = 0.00015; Figure 1K) and CTCF (p = 1.35e-12; Figure 1L) binding sites. Although enrichment scores were less robust, regions losing RD with age were significantly enriched in genes (p = 8.725e-05; Figure 1G), promoters (p = 7.073e-11; Figure 1H), and PRC2 target genes (p = 0.020; Figure 1J) and were depleted in enhancers (p = 0.030; Figure 1I). The mean CpG density did not differ in age-associated regions when compared to background (Figure 1M). Genes which accumulate disorder with age (n = 1,635) were significantly enriched in 552 different biological processes (GO:BP) with the most significant terms relating to nervous system development and differentiation (Table S1). Genes losing disorder with age (n = 197) were enriched in 14 different biological processes, with the most significant terms relating to multicellular organismal development (Table S2).

We next examined the relationship between RD and regional averages of Shannon’s Entropy, a commonly used measure of epigenetic drift. Regional entropy (RE) is calculated directly from mean methylation values, and thus has a strong relationship to regional methylation (RM), even when regional values are averaged across all individuals (Figure 2A). However, in loci where RE reaches its maximum (RE = 1, mean methylation = 50%), RD spans from fully ordered to fully disordered (RD = 0-1). We found that the relationship between average RD and average RE is best explained by a quadratic relationship (R^2^ = 0.83, p < 2.2e-16; Figure 2B) with increasing RE generally indicating increases in RD. While the relationship between global RD and global RE does not change with age (Figure S2), the relationship becomes increasingly variable at greater values of RE (Figure S3). Age dependent changes to RM and RD are linked as 78.7% (n = 38,926) of regions experiencing modest age-associated RD (cor >= |0.25|) also incur modest age-associated RM (cor >= |0.25|); however, the remaining 21.3% (n = 10,531) of age-associated changes in RD do not correspond with RM, and 34.1% (n = 20,152) of age-associated RM occur independently of changes in RD (Figure 2C).

**Figure 2.**
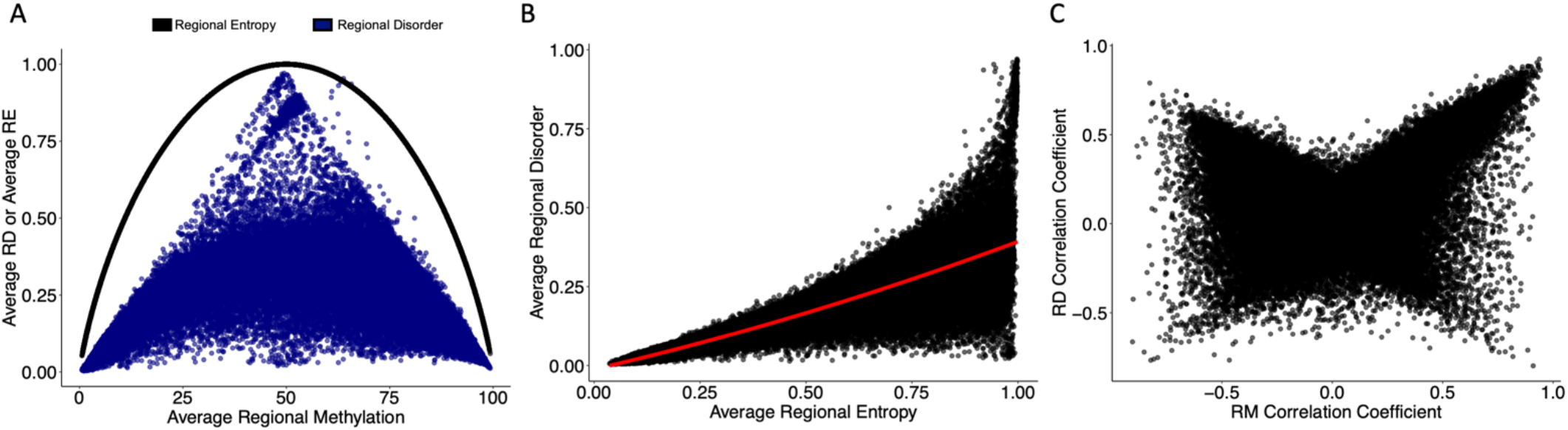
Regional disorder is distinct from Shannon’s entropy and age-associated changes in mean methylation. (A) The relationship of regional entropy (RE; black line) with regional methylation and regional disorder (RD; blue dots) with regional methylation (RM). Data points show a single region averaged across all samples. (B) Relationship between RD and RE averaged across all samples. (C) Correlation coefficients of RD with age and RM with age across the 153 samples used to build the epigenetic clock. Regions which increase in RM or RD with age have positive correlation coefficients, regions which decrease in RM or RD with age have negative correlation coefficients.

We also aimed to understand how signals underlying epigenetic clocks relate to epigenetic disorder. Interestingly, there is a clear enrichment of Petkovich epigenetic clock loci in regions which increase and decrease in RD with age (Figure 3A), with the absolute correlation coefficient of RD and age being significantly higher in Petkovich epigenetic clock regions when compared to those not included in the clock (p < 2e-16; Figure 3B). However, 37 of the 90 total clock CpGs fall into the same 200 bp genomic region. To more thoroughly resolve the relationship between epigenetic clock signals and epigenetic disorder, we built a series of epigenetic clocks based on CpG, RM, RE, and RD states. Over the 153 LOO iterations for each clock type, there was no difference in absolute error across clocks, suggesting that each methylation metric is capable of predicting chronological age with equivalent accuracy (Figure 3C). Similarly, there was no difference in the mean absolute error produced by the representative clocks (Figure S1).

**Figure 3.**
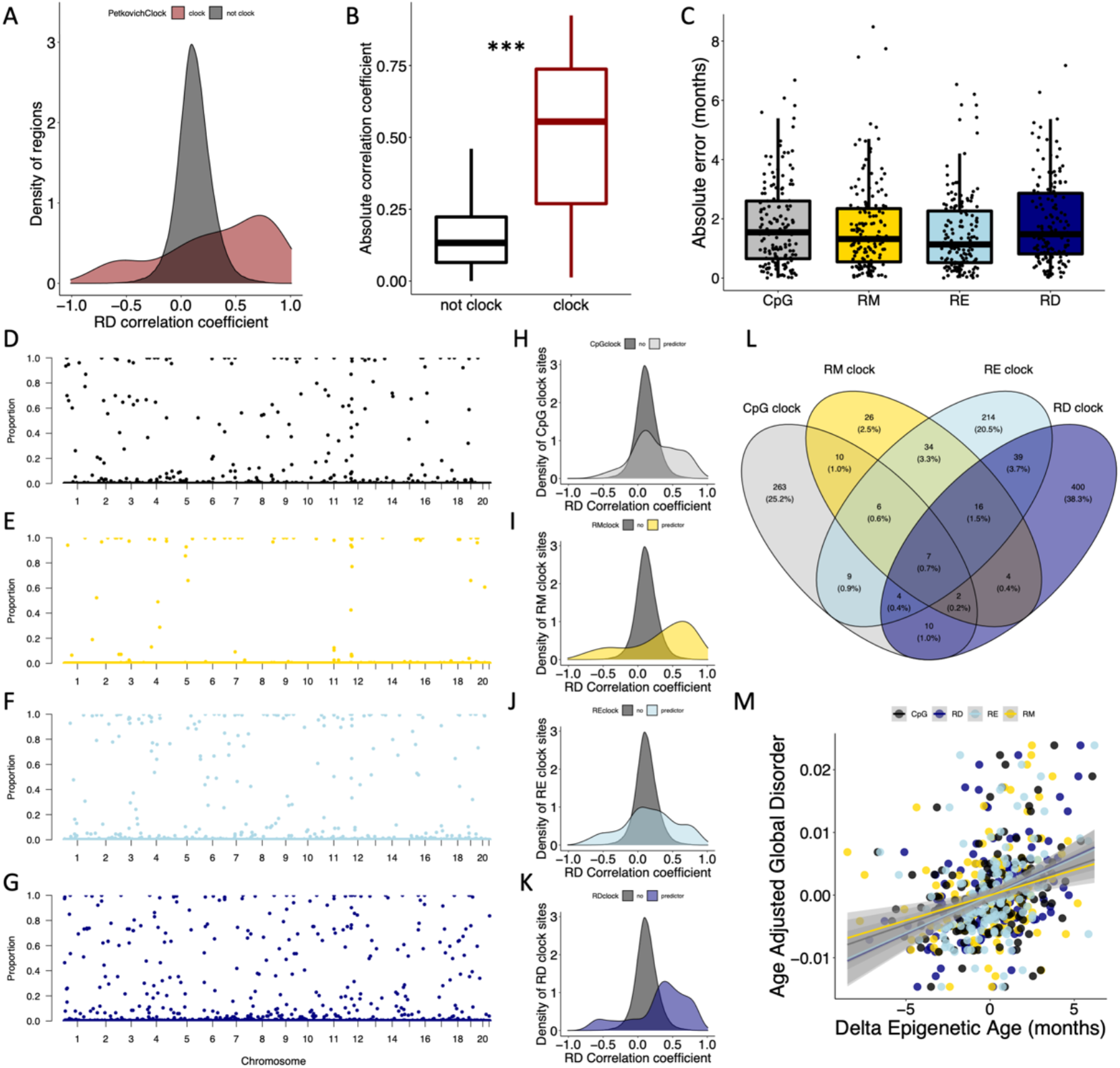
Epigenetic disorder underlies epigenetic clock signals. (A) Distribution of Petkovich epigenetic clock sites (red) across correlation coefficients between regional disorder (RD) and age. (B) Average absolute correlation coefficient between RD and age of regions which are included in the Petkovich epigenetic clock (red) compared to those which are not included. (C) Error of epigenetic age estimates produced by leave-one-out cross validation (LOOCV) for each data type. (D-G) Manhattan plots showing the robustness for each region (i.e., the proportion of clocks each region was selected in) across (D) CpG methylation (black), (E) regional methylation (RM; yellow), (F) regional entropy (RE; light blue), and (G) RD (dark blue) contexts. (H-K) Density plots showing the distribution of clock sites for each data type across correlation coefficients between RD and age. (L) Overlap between regions included in epigenetic clocks produced from each data type. (M) Relationship between delta epigenetic age (chronological age – predicted age) and age-adjusted global disorder for each data type.

To further compare the influence of methylation context on clock composition, we assessed the overlap of loci incorporated into each clock type as well as the frequency in which they were selected (referred to as robustness). Of the LOO iterations, the CpG clocks selected 312 different regions with an average robustness of 0.11 (Figure 3D), RM clocks selected 106 different regions with an average robustness of 0.05 (Figure 3E), RE clocks selected 330 different regions with an average robustness of 0.11 (Figure 3F), and RD clocks selected 483 different regions with an average robustness of 0.13 (Figure 3G). Interestingly, the mean absolute RD correlation coefficients for age were significantly higher for CpG, RM, RE and RD clock regions when compared to nonclock regions (CpG p < 2e-16, RM p < 2e-16, RE p < 2e-16, RD p < 2e-16; Figure 3H-K). The majority of clock sites (86.5%) were specific to each clock type; however, seven regions were selected across all clock types. Pan-clock regions are all associated with genes (Map10, Nlrp5-ps, Rasef, Rnf220, Evx2, Gm21297, and Apba1), with five (71%) regions located within promoters, and all increase in disorder with age (Figure 3L). While all four datatypes produce low errors in age prediction (Figure 3C), the discordance of chronological age with the age prediction (or “delta epigenetic age”) is highly correlated with age adjusted global disorder in all datatypes (CpG R^2^ = 0.09, p = 7.53e-05; RM R^2^ = 0.07, p = 0.0007, RE R^2^ = 0.12, p = 6.89e-06, RD R^2^ = 0.18; p = 3.00e-08; Figure 3M).

We then tested the influence of common lifespan manipulations on epigenetic age estimates across different clock types. Caloric restriction led to a reduction in age-associated RD, but the effect varied across strains. Male C57BL/6 mice fed a calorie restricted diet had significantly younger epigenetic ages when compared to controls as determined by all clock types (CpG p = 0.00043, RM p = 2.16e-07, RE p = 4.62e-06, RD p = 5.75e-08; Figure 4A). However, mean age adjusted global disorder appeared unaffected (p = 0.62; Figure 4B). Conversely, male B6D2F1 mice fed a calorie restricted diet only had significantly younger epigenetic ages as determined by the RM epigenetic clock (CpG p = 0.46, RM p = 0.018, RE p = 0.073, RD p = 0.39; Figure 4C). However, there was a slight trend for calorie restricted individuals to have greater mean age adjusted global disorder when compared to mice on a standard diet (p = 0.097; Figure 4D). Genetic interventions which extend lifespan resulted in a general decrease in epigenetic age. Snell Dwarf mice had significantly younger epigenetic ages when compared to controls according to the RD and RM clocks, but not the CpG or RE clocks (CpG p = 0.078, RM p = 0.0059, RE p = 0.23, RD p = 0.011; Figure 4E). Snell Dwarf mice also showed reduced mean age adjusted global disorder compared to control mice (p = 0.027; Figure 4F). GHR knock out also resulted in significantly younger epigenetic ages according to the RM and RE clocks, but not the CpG or RD clocks (CpG p = 0.088, RM p = 0.0080, RE p = 0.031, RD p = 0.64; Figure 4G), and no difference in age adjusted regional disorder was observed between GHRKO and control mice (p = 0.39; Figure 4H).

**Figure 4.**
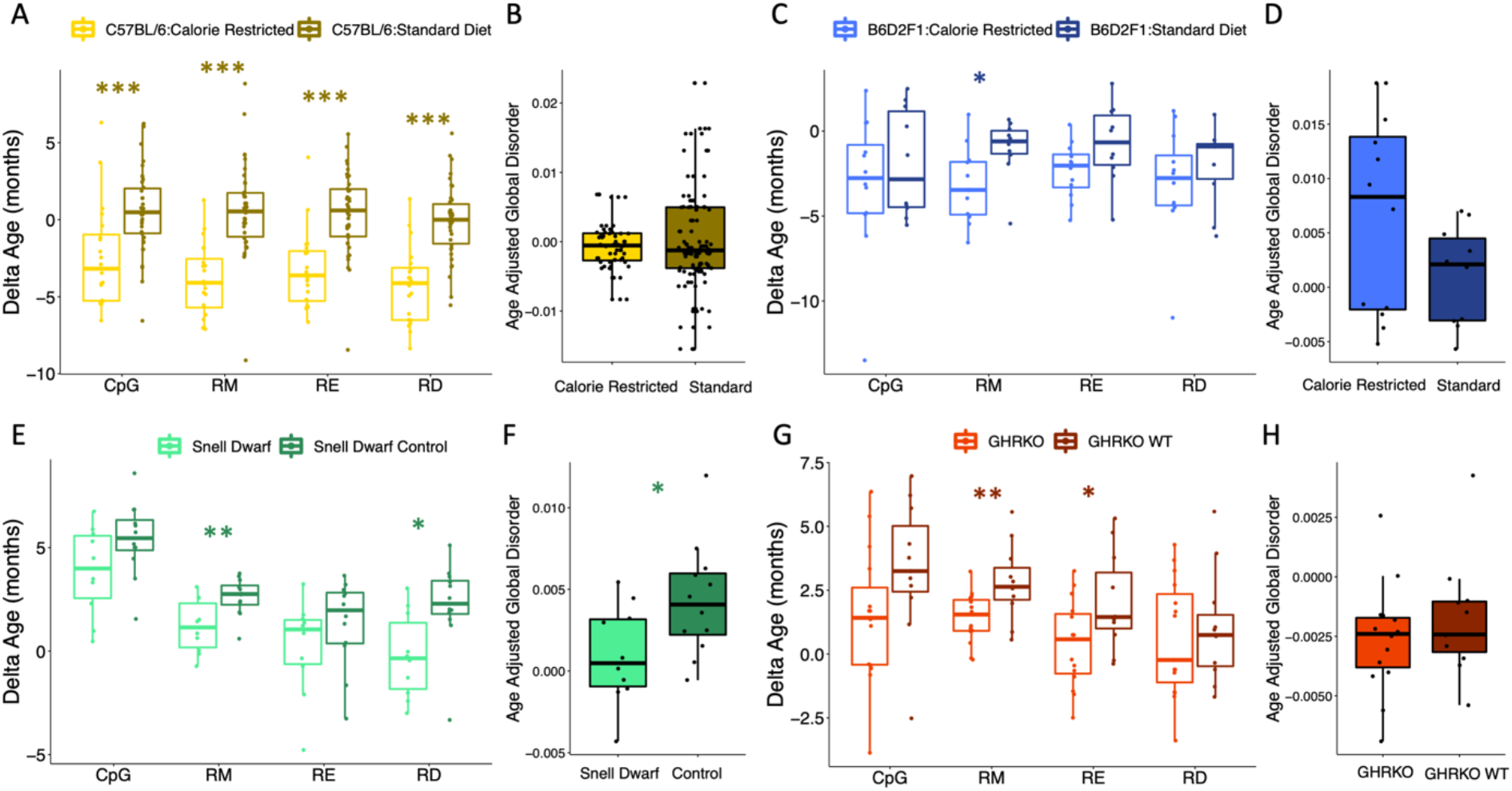
Epigenetic disorder is influenced by lifespan extending manipulations. The effect of caloric restriction in C57BL/6 mice on (A) epigenetic age predictions from each data type and (B) age-adjusted global disorder. The effect of caloric restriction in B6D2F1 mice on (C) epigenetic age predictions from each data type and (D) age-adjusted global disorder. Comparison of Snell dwarf and control mice on (E) epigenetic age predictions from each data type and (G) age-adjusted global disorder. The effect of growth hormone receptor knock-out (GHRKO) on (G) epigenetic age predictions from each data type and (H) age-adjusted global disorder. All plots show median, upper, and lower quartiles, and maximum and minimum. Outliers beyond 1.5 interquartile range are plotted.

We next investigated the impacts of cellular dedifferentiation on epigenetic disorder by comparing DNA methylomes of iPSC cells and their differentiated precursors. A significant reduction in epigenetic age predictions after dedifferentiation was observed across all clock types except for RE (CpG p = 7.11e-08, RM p = 3.41e-06, RE p = 0.20, RD p = 3.93e-08; Figure 5A). Interestingly, there was no difference in the global disorder between kidney or lung fibroblasts when compared to their respective iPSCs (p = 0.28; Figure 5B). Given that dedifferentiation led to a reduction in epigenetic age estimates but did not affect global disorder, we sought to identify those regions in which RD is affected by dedifferentiation. Upon comparing RD across all fibroblasts and iPSCs, 26,512 regions significantly increase in RD after differentiation and 19,419 regions significantly decrease in disorder after dedifferentiation, but these regions do not disproportionately acquire age-associated RD relative to background (Figure 5C). Interestingly, the influence of dedifferentiation on RD epigenetic clock estimates are driven by differences in RD at just several clock sites (Figure 5D), with four regions contributing 35.7% of the overall effect.

**Figure 5.**
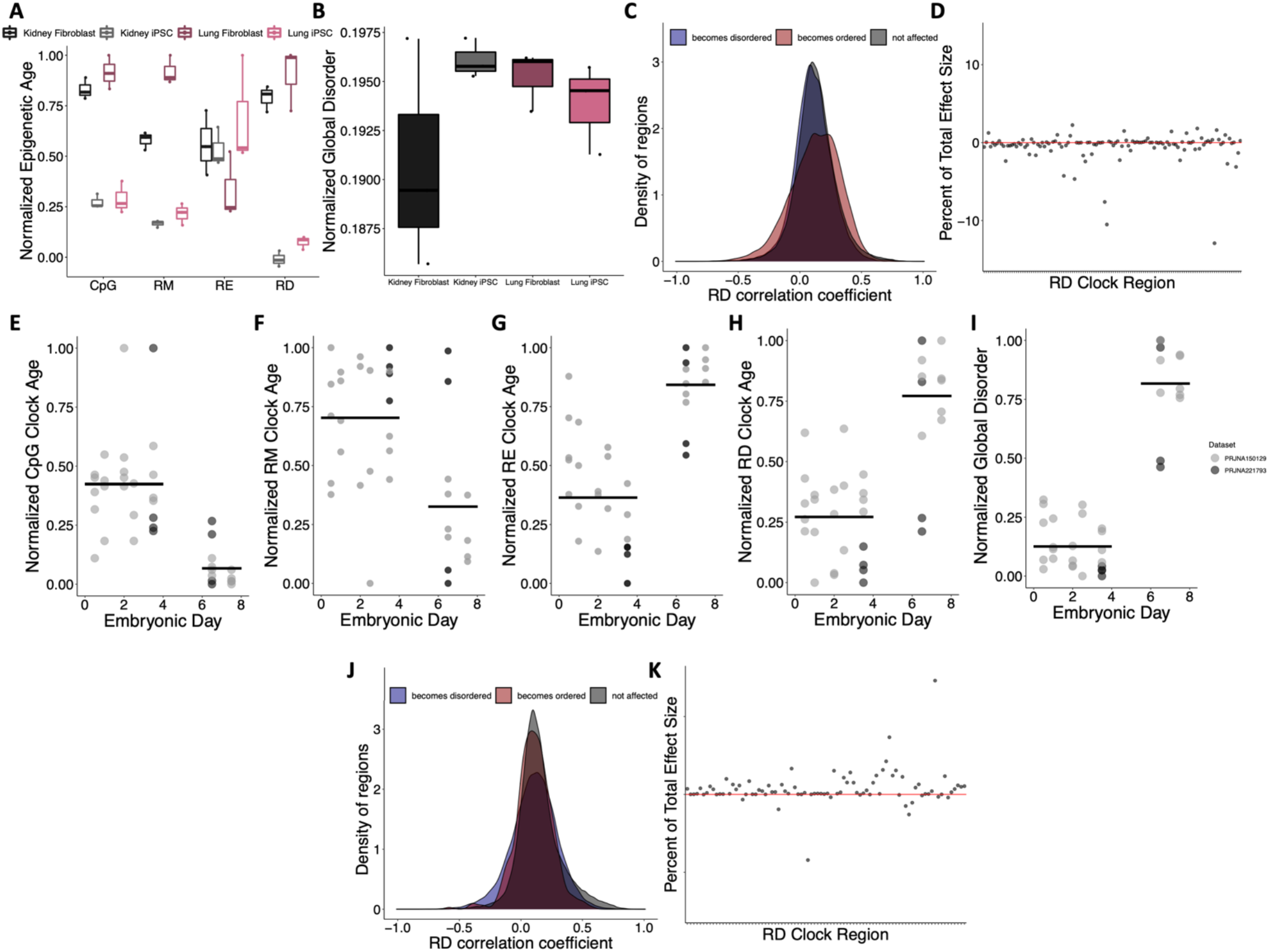
Epigenetic disorder during de-differentiation and development. (A) Epigenetic age predictions using each of the representative epigenetic clocks and (B) global disorder of kidney fibroblasts (black), kidney derived iPSCs (grey), lung fibroblasts (dark purple), and lung derived iPSCs (pink). Plot shows median, upper and lower quartiles, maximum, and minimum. Outliers beyond 1.5 interquartile range are plotted. (C) Distribution of regions which gain (blue) or lose (red) disorder after de-differentiation across correlation coefficients between regional disorder (RD) and age. (D) Effect sizes of dedifferentiation on the RD epigenetic clock. Stubbs CpG methylation (E), RM (F), RE (G), and RD (H) epigenetic clock predictions of samples during embryonic development. (I) Global disorder of samples during embryonic development. (J) distribution of regions which gain (blue) or lose (red) disorder during early development across correlation coefficients between regional disorder (RD) and age. (K) Effect sizes of development on the RD epigenetic clock.

Consistent with a previous report identifying an epigenetic rejuvenation event occurring during early development [16], we observed a significant decrease in epigenetic age predictions occurring between embryonic days 4 and 6 using the Stubbs CpG methylation epigenetic clock [7] (CpG p = 1.32e-08; Figure 5E) and the RM clock (RM p = 0.0019; Figure 5F). Conversely, we observe a significant increase in epigenetic age predictions during early embryonic development when using the RD and RE clocks (RD p = 3.81e- 05; RE p = 1.90e-08; Figure 5G-H). In addition, global disorder is strongly increased during development (Figure 5I). Upon comparing RD across all samples from embryonic days 0.5-3.5 (n = 24) and embryonic days 6.5-7.5 (n = 12), 31,687 regions significantly increase in RD during development and 368 regions significantly decrease in RD during development. These regions were not significantly enriched in regions with age-associated RD (Figure 5J). Similar to our findings examining the influence of de-differentiation, the predictions of the epigenetic clock appear to be driven strongly by differences in RD at just several clock sites, with one region having a total effect size of 2.51 months, contributing 14.6% of the difference in ages between groups (Figure 5K). Given that the clock has 86 of the regions included as predictors represented, we would expect each region to contribute just 1.16% to the overall effect size.

## Discussion

Epigenetic drift is broadly hypothesized to be a primary contributor to epigenetic aging. However, drift is a multifaceted phenomenon encompassing both stochastic and deterministic processes, and is unlikely to be fully captured by a single metric. In this study, we report an approach for spatially resolving genomic patterns of DNA methylation disorder, which is distinct from measures of both average methylation and entropy. Age-associated changes in regional disorder (RD) are found in approximately one third of the genome and generally reflect the accumulation of disordered methylation; however, the opposite pattern is observed in a subset of regions in which DNA methylation patterns become more ordered with age. Given that epigenetic drift or disorder is thought to be driven in part by stochastic processes, we hypothesize that the directionality of changes in RD represent different functional pathways. Yet, the specific biological mechanisms that mediate losses of RD with age remain unclear. Age-associated gains and losses of RD were disproportionately observed in coding regions, promoters, and regions harboring PRC2 target genes, and age-associated increases in RD were strongly enriched in developmental genes, especially those functioning in neural development. These gains in disorder with age support the deleteriome model of aging [42], wherein small deleterious errors accumulate in the epigenome without effect until later in life, when epigenomic stability is compromised [2]. We suggest that disorder accumulates across the genome until it reaches a critical threshold – this may explain why the majority of regions across the genome are characterized by relatively low disorder (RD < 0.5; data not shown). The value of this hypothetical threshold and the factors which contribute to the accumulation of disorder have the potential to explain the rate of aging and possibly maximum lifespan across species [36].

Consistent with the hypothesis that disorder in DNA methylation patterns underlies signals in conventional CpG clocks (i.e., those based on mean CpG methylation levels), we find that loci comprising clocks constructed using RD, RM, RE, and CpG contexts are all enriched for regions in which disorder changes with age. However, we identify notable distinctions and minimal overlap across contexts. For example, while a subset of CpGs selected as predictors in an epigenetic clock were enriched in regions with age-associated disorder, many CpG clock sites also fell into regions lacking age-associated changes in disorder. Thus, while disorder underlies some components of traditional CpG epigenetic clocks, other components may be attributed to other processes like coordinated changes in methylation or cell type composition. By contrasting the effects of caloric restriction, genetic manipulations, cellular reprogramming, and development across different clock types, we further identify both similarities and clear distinctions according to DNA methylation context and genomic scale. For example, while traditional lifespan extending treatments, such as caloric restriction, broadly affect RD epigenetic clocks, there is no observable effects on global disorder. Similarly, while CpG and RM clocks demonstrate a “ground zero” occurring during mid-development [16], we see an increase in RE and RD clock predicted ages during the same period. Collectively, these findings demonstrate the connections between epigenetic drift and other aspects of epigenetic aging, while also highlighting a complexity that should be considered when assessing read outs from epigenetic clocks alone.

In mice, global disorder changes with age according to a quadratic function, with decreases in disorder occurring rapidly earlier in life prior to a steady increase with age. This pattern is consistent with previous findings of a quadratic relationship between global DNA methylation entropy and age in the naked-mole rat [43]. The initial high level of global disorder suggests that development, as well as aging, may be characterized by a disorganized epigenetic landscape – possibly due to a transitionary period between methylation states. Given the dynamic nature of the DNA methylome during development [6,44,45], it is likely that RD metrics, like other measures of DNA methylation that provide temporal snapshots, capture this transition as high disorder. While data from embryonic samples suggest that disorder increases during early development, the trajectory of global disorder throughout development, and whether it corresponds with previous findings of an epigenetic “ground zero” during development [16], will require a more complete developmental series to fully resolve.

Age estimates derived from epigenetic clocks are ideal for predicting chronological age (i.e., forensics, conservation and management applications [46,47]) as well as identifying the consequences of accelerated epigenetic aging (i.e., biomarkers in biomedical approaches [1]). Yet, collapsing mean methylation levels into a single value presents challenges for understanding the drivers and biological pathways responsible for epigenetic aging. Given the push towards targeted, highthroughput approaches (e.g., bead-based assays) for acquiring data on age-associated methylation [48,49], critical biological information is missed. While CpG level resolution has been integral in developing our understanding of epigenetic aging, clocks built using regionally averaged methylation perform with similar accuracy to those trained on individual CpGs. We further demonstrate that the effect size of individual clock sites varies widely, and thus, changes in methylation states of just one or several clock loci can be misinterpreted as wholesale changes in epigenetic age. This is important especially when age estimates are compared across studies and different datasets. For example, we report that a single RD clock region accounted for nearly 10% of the difference in age estimation between fibroblasts and iPSCs. While the age predictions generated corroborate previous findings [6,40], the inclusion (or exclusion) of this single region vastly changes our interpretation of the effects of de-differentiation on epigenetic age. Thus, using epigenetic clocks of any kind gives us a narrow, and potentially easily skewed, understanding of epigenetic aging at the genomic scale.

Overall, this study provides robust empirical evidence that epigenetic drift, as measured by epigenetic disorder, accumulates with age in non-random places of the mouse genome. Our analyses suggest that epigenetic disorder underlies aspects of traditional epigenetic clocks and highlights critical gaps in our interpretation of epigenetic aging. Although more work needs to be done to better resolve the drivers of epigenetic disorder – we provide an empirical basis for testing assumptions about this emerging phenomenon.

## Acknowledgements

This material is based upon work supported by the Department of Energy Office of Environmental Management under Award Number DE-EM0005228 to the University of Georgia Research Foundation. In addition, this work was supported by awards from the National Science Foundation (Award No. 2026210) and the National Institutes of Health (Award No. 1R56AG078336-01).

## Disclaimer

This report was prepared as an account of work sponsored by an agency of the United States Government. Neither the United States Government nor any agency thereof, nor any of their employees, makes any warranty, express or implied, or assumes any legal liability or responsibility for the accuracy, completeness. Or usefulness of any information, apparatus, product, or process disclosed, or represents that its use would not infringe privately owned rights. Reference herein to any specific commercial product, process, or service by trade name, trademark, manufacturer, or otherwise does not necessarily constitute or imply its endorsement, recommendation, or favoring by the United States.

## Supplemental Material

**Figure S1:**
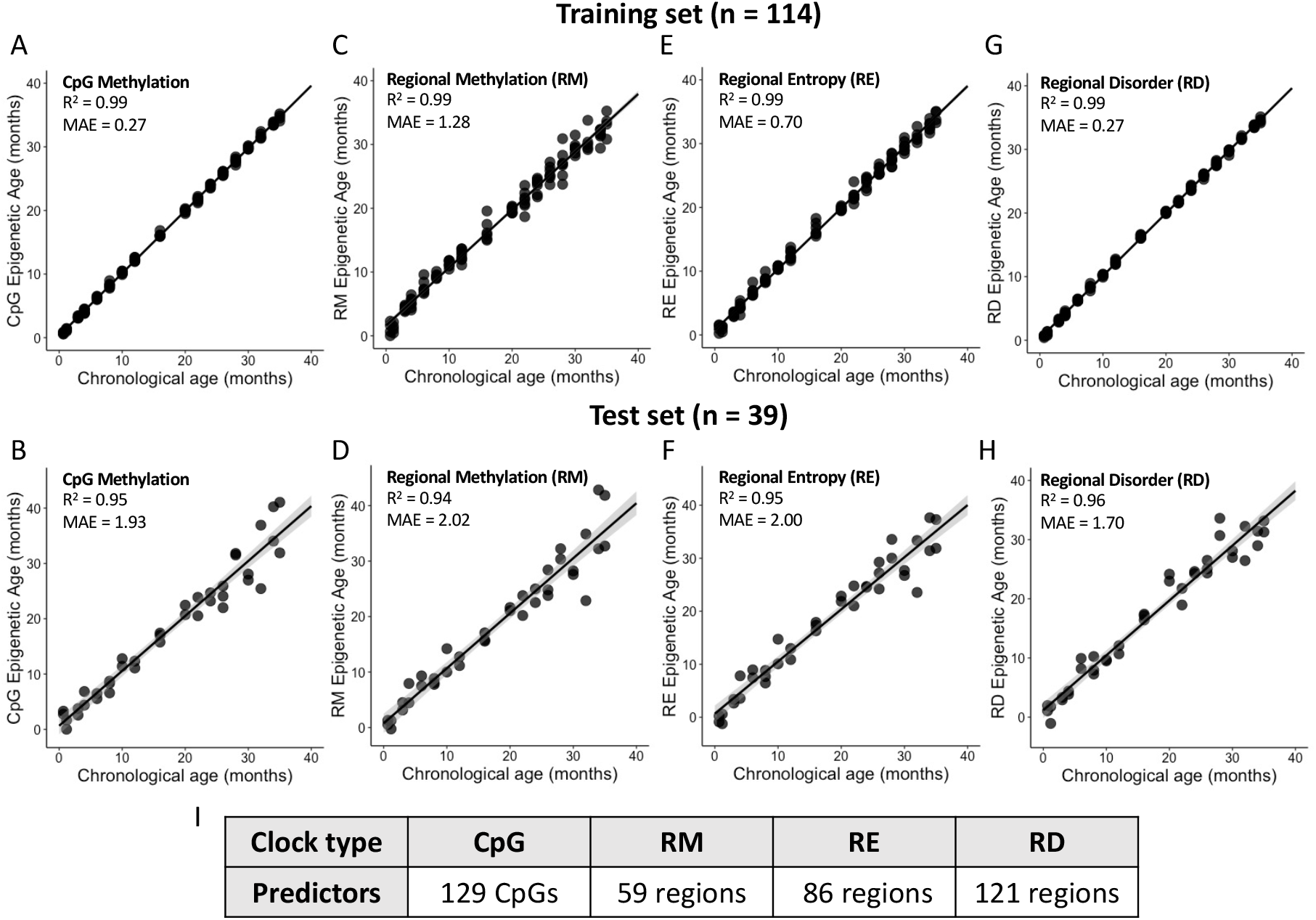
Representative epigenetic clocks from each datatype. (A) and (B) show results for the epigenetic clock constructed using CpG methylation data, (C) and (D) show results for the epigenetic clock constructed using regional methylation (RM) data, (E) and (F) show results from an epigenetic clock based on regional entropy (RE) data, and (G) and (H) show results from the epigenetic clock based on regional disorder (RD) data. Panel (I) indicates the number of predictors comprising each clock.

**Figure S2:**
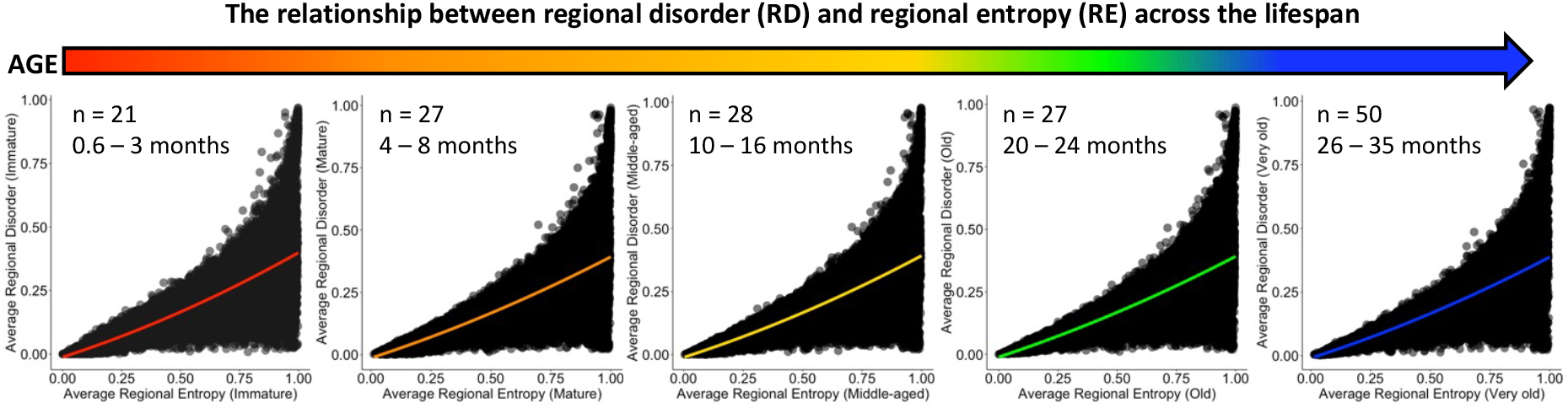
The relationship between average regional disorder and average regional entropy across the lifespan of mice. Samples are broken up into distinct life stages which are denoted by different colors (immature – red, mature – orange, middleaged – yellow, old – green, and very old – blue).

**Figure S3:**
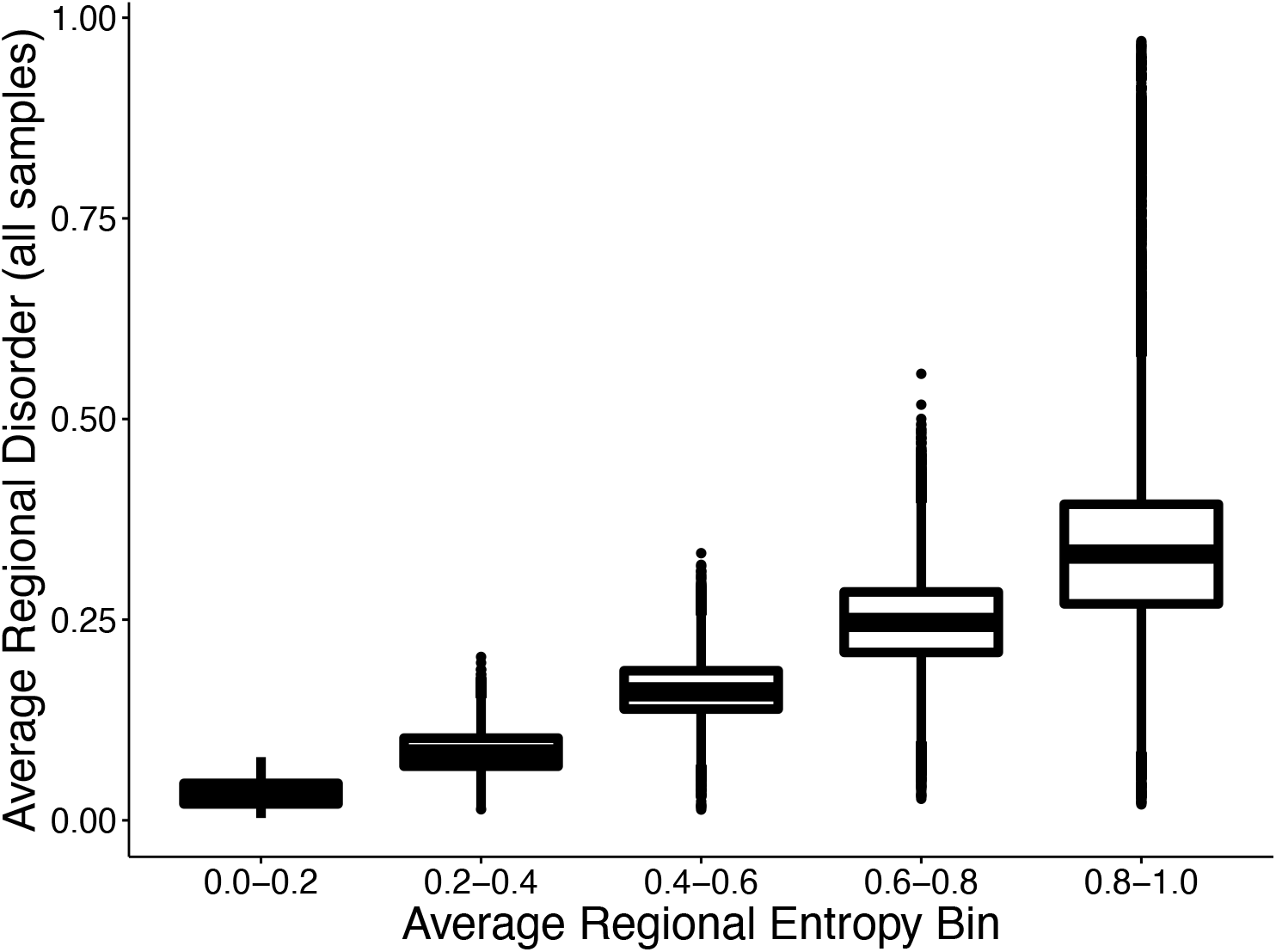
Variation in average regional disorder increases with increasing values of average regional entropy.

**Table S1:** Gene ontology results from regions with age-associated gains in disorder. Top 20 terms from molecular function (MF) and biological process (BP) based on significance values are shown.

| Source | GO Term Name | GO ID | Adjusted p-value |
| --- | --- | --- | --- |
| GO:MF | DNA-binding transcription factor activity, RNA polymerase II-specific | GO:0000981 | 1.76E-28 |
| GO:MF | DNA-binding transcription factor activity | GO:0003700 | 7.58E-28 |
| GO:MF | sequence-specific double-stranded DNA binding | GO:1990837 | 5.72E-23 |
| GO:MF | double-stranded DNA binding | GO:0003690 | 4.64E-21 |
| GO:MF | sequence-specific DNA binding | GO:0043565 | 1.37E-20 |
| GO:MF | RNA polymerase II transcription regulatory region sequence-specific DNA binding | GO:0000977 | 1.79E-20 |
| GO:MF | transcription regulator activity | GO:0140110 | 5.41E-20 |
| GO:MF | transcription cis-regulatory region binding | GO:0000976 | 7.85E-19 |
| GO:MF | transcription regulatory region nucleic acid binding | GO:0001067 | 1.17E-18 |
| GO:MF | RNA polymerase II cis-regulatory region sequence-specific DNA binding | GO:0000978 | 1.70E-17 |
| GO:MF | cis-regulatory region sequence-specific DNA binding | GO:0000987 | 2.57E-17 |
| GO:MF | binding | GO:0005488 | 8.47E-17 |
| GO:MF | protein binding | GO:0005515 | 1.12E-15 |
| GO:MF | DNA-binding transcription activator activity | GO:0001216 | 1.01E-11 |
| GO:MF | DNA-binding transcription activator activity, RNA polymerase II-specific | GO:0001228 | 1.31E-11 |
| GO:MF | gated channel activity | GO:0022836 | 1.80E-11 |
| GO:MF | DNA binding | GO:0003677 | 8.54E-09 |
| GO:MF | ion channel activity | GO:0005216 | 8.80E-09 |
| GO:MF | voltage-gated cation channel activity | GO:0022843 | 1.21E-08 |
| GO:MF | channel activity | GO:0015267 | 2.18E-08 |
| GO:BP | nervous system development | GO:0007399 | 9.36E-84 |
| GO:BP | neurogenesis | GO:0022008 | 1.71E-68 |
| GO:BP | system development | GO:0048731 | 4.56E-67 |
| GO:BP | multicellular organism development | GO:0007275 | 3.14E-65 |
| GO:BP | generation of neurons | GO:0048699 | 4.99E-63 |
| GO:BP | anatomical structure development | GO:0048856 | 4.01E-61 |
| GO:BP | neuron differentiation | GO:0030182 | 3.05E-60 |
| GO:BP | multicellular organismal process | GO:0032501 | 5.25E-58 |
| GO:BP | developmental process | GO:0032502 | 5.17E-57 |
| GO:BP | anatomical structure morphogenesis | GO:0009653 | 1.15E-54 |
| GO:BP | cell-cell signaling | GO:0007267 | 1.09E-50 |
| GO:BP | central nervous system development | GO:0007417 | 4.10E-48 |
| GO:BP | neuron development | GO:0048666 | 2.56E-47 |
| GO:BP | cell differentiation | GO:0030154 | 5.75E-45 |
| GO:BP | cell development | GO:0048468 | 9.25E-45 |
| GO:BP | cellular developmental process | GO:0048869 | 1.71E-44 |
| GO:BP | neuron projection development | GO:0031175 | 2.60E-41 |
| GO:BP | animal organ development | GO:0048513 | 3.67E-41 |
| GO:BP | brain development | GO:0007420 | 2.06E-40 |
| GO:BP | head development | GO:0060322 | 4.07E-40 |

**Table 5.S2:**
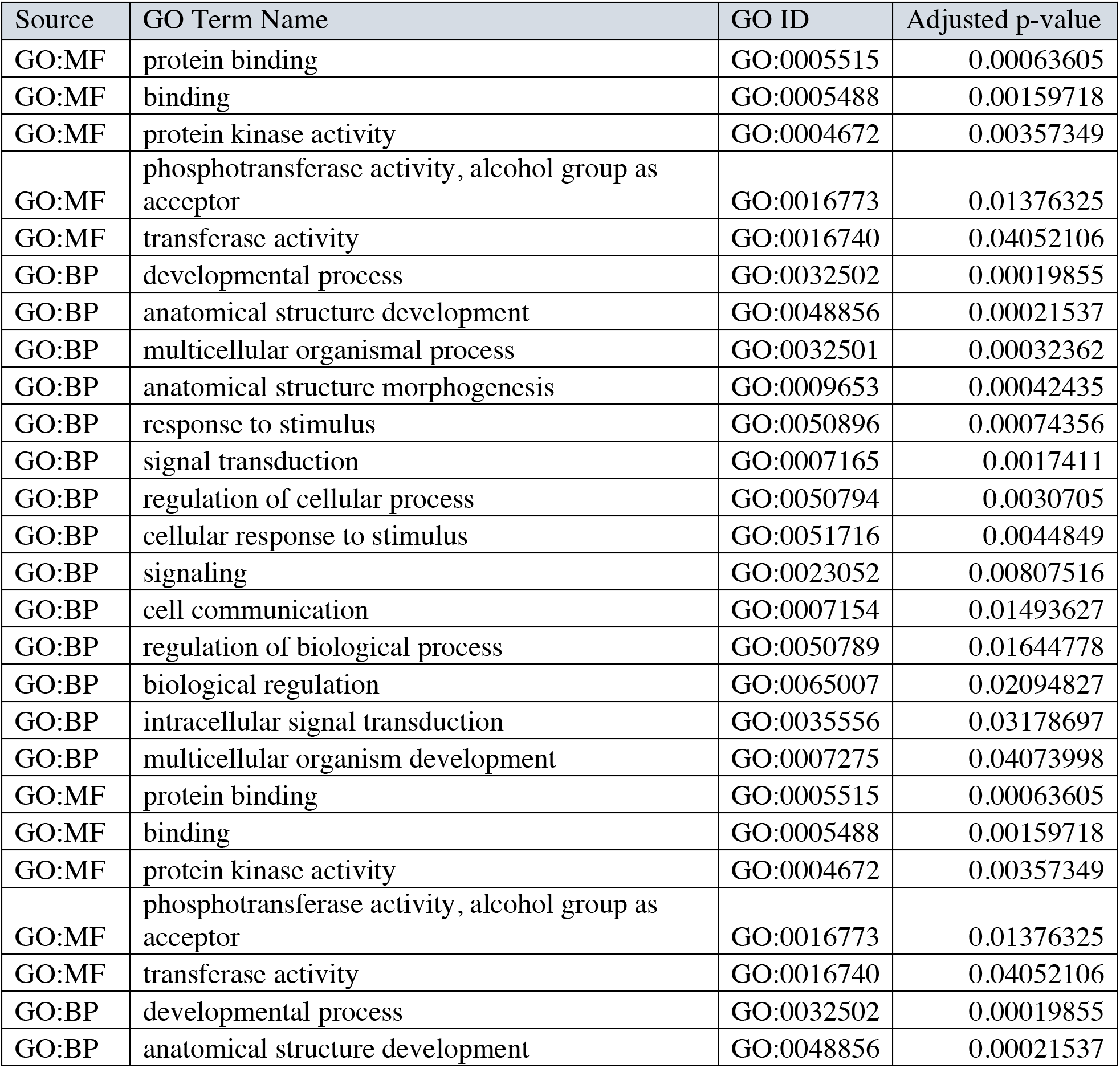
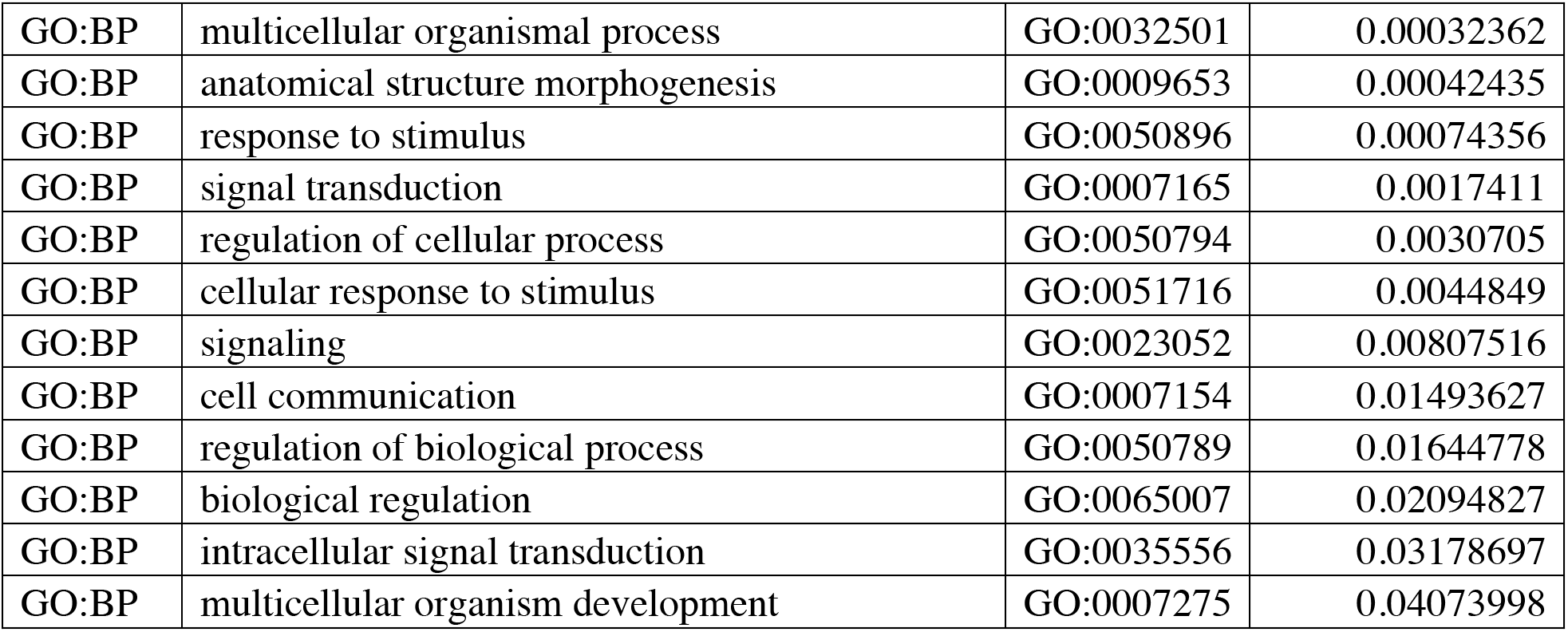
Gene ontology results from regions with age-associated losses in disorder. Terms from molecular function (MF) and biological process (BP) are shown.

